# Dynamics of PAR proteins explain the oscillation and ratcheting mechanisms in dorsal closure

**DOI:** 10.1101/348540

**Authors:** C.H. Durney, T.J.C. Harris, J.J. Feng

## Abstract

We present a vertex-based model for *Drosophila* dorsal closure that predicts the mechanics of cell oscillation and contraction from the dynamics of the PAR proteins. Based on experimental observations of how aPKC, Par-6 and Bazooka migrate from the circumference of the apical surface to the medial domain, and how they interact with each other and ultimately regulate the apicomedial actomyosin, we formulate a system of differential equations that capture the key features of the process. The oscillation in cell area in the early phase of dorsal closure results from an intracellular negative feedback loop that involves myosin, an actomyosin regulator, aPKC and Bazooka. In the slow phase, gradual sequestration of apicomedial aPKC into Bazooka clusters causes incomplete disassembly of the myosin network over each cycle of oscillation, thus producing the so-called ratchet. The fast phase of rapid cell and tissue contraction arises when medial myosin, no longer hindered by aPKC, builds up in time and produces sustained contraction. Thus, a minimal set of rules governing the dynamics of the PAR proteins, extracted from experimental observations, can account for all major mechanical outcomes of dorsal closure, including the transitions between its three distinct phases.

Insert Received for publication Date and in final form Date

## INTRODUCTION

Dorsal closure (DC) is an important morphogenetic process during the embryonic development of *Drosophila.* During DC, two flanking epidermal tissues are fused together to close an opening on the dorsal side of the embryo, covering up the thin amnioserosa (AS) tissue. The entire DC process, lasting about 3.5 hours, exhibits three distinct phases showing great spatial and temporal coordination. The early phase is characterized by AS cell oscillations and no loss in overall tissue area. The ensuing slow phase exhibits a gradual loss in overall tissue area and a dampening of cellular oscillations. Finally, the fast phase displays rapid and persistent loss of tissue area and cessation of the oscillations. While recent experiments have identified several contributing factors in dorsal closure (1–3), the connections between them remain elusive.

For the early phase, the key question is the cause of the cyclic oscillation of the cell area. It has been observed that on the apical surface, a medial network of actomyosin periodically assembles and disassembles, with a period of approximately 4 minutes (1, 4–6). The cell contracts following the assembly of the actomyosin network, and relaxes when the network disassembles (4, 7). After each cycle, the cell area returns to more or less the previous state. The contraction of one cell pulls on its neighbors, yielding a preference for antiphase pulsing between neighboring cells (1). But Blanchard et al. (3) have also observed transient “ribbons” of cells that are simultaneously contracting or expanding. It remains to be ascertained whether the oscillatory behavior originates from the autonomous dynamics within each cell or from the communications between neighbors, or even both (1, 8, 9).

Naturally one seeks an explanation for the oscillatory behavior from biochemical signaling (4, 10–13). David et al. (4) highlighted the roles of the PAR proteins. Bazooka (Baz) promotes the duration of actomyosin contraction, while the Par-6/aPKC complex promotes the lull time between contractions. More recently, David et al. (12) further delineated the spatial and temporal relationship among the PAR proteins and the actomyosin network that periodically assembles in the apicomedial domain of the AS cells. The actomyosin recruits aPKC from the circumference of the apical surface toward the medial region, where it forms puncta that colocalize with the actomyosin network. In turn, aPKC recruits Bazooka into the medial region. As aPKC is known to inhibit actomyosin, and the formation and stabilization of aPKC-Baz complexes result in apical constriction, David et al. (12) hypothesized that the PAR proteins form a negative feedback loop with the medial actomyosin network to produce the oscillatory outcome. This is the strongest evidence so far in favor of a cell-autonomous cause of the oscillation.

To rationalize the areal contraction of the slow phase, the two central ideas are the ratcheting mechanism and the supracellular actin cable, although recent evidence has also suggested cell volume loss as a contributor (14). The concept of an intracellular or internal ratchet was introduced by Martin et al. (7, 15) to explain the pulsed constriction of *Drosophila* ventral furrow cells. After each contraction, a cell is prevented from returning to its prior apical area by tension emanating from an apicomedial actomyosin meshwork, which stabilizes the cell shape and effect net constriction. Consistent with the above, the live images of David et al. (4, 12) demonstrate an apicomedial actomyosin network that grows stronger over each pulse in the slow phase, a transition that coincides with greater aPKC-Baz interaction in the medial domain. Roughly concurrent with the onset of the slow phase, an actin cable appears along the boundary between the AS tissue and the surrounding epidermis. It encircles the AS like a purse string and clamps down on the tissue. This was initially considered the main cause of AS contraction (1, 16), but more recent work has challenged its importance (6, 15, 17–19). For example, Wells et al. (6) disrupted the purse string by severing one or both of the canthi and observed that closure proceeded normally. Pasakarnis et al. (18) used selective force elimination to demonstrate that the amnioserosa cells alone are capable of driving dorsal closure. Ducuing and Vincent (19) reported that the actin cable does not contribute appreciably to the dorsal-ward movement of the leading edges. Thus, a consensus seems to be forming on a secondary role for the actin cable.

The advent of the fast phase brings a few new factors. As the AS tissue narrows, the opposing epidermis extends filopodia onto the opposite side, which then pull to help the closure (20–22). This is known as zippering. Much like the actin cable, the zippering forces have been shown to be superfluous for complete closure (6). Moreover, AS cell apoptosis contributes to rapid closure by dropping out of the plane of the amnioserosa, effectively reducing the tissue area (2, 5, 16, 23). Finally, the epidermis may elongate actively to assist rapid closure in the fast phase (24).

Mathematical modeling has been used to integrate the experimental insights above into a more precise and quantitative understanding (21). A majority of these models are purely mechanical, imposing prescribed forces on the tissue (25) or on individual cells (1, 26) and examining their elastic or viscoelastic responses (27, 28). More recent work strove to integrate the mechanical and biochemical aspects of the process (29, 30). Wang et al. (17) assumed a simple kinetic equation for a hypothetical signaling molecule, and predicted cyclic myosin dynamics for the early phase from a negative feedback loop. By an *ad hoc* device of shrinking the resting length of the elastic cell borders, their model produced net loss of cell and tissue area. Similarly, Machado et al. (8) relied on a feedback loop between a putative signaling molecule and actin turnover, which was a proxy for myosin attachment and detachment. This model showed how the rate of actin turnover may modulate the period of AS cell oscillation. These models are commendable in seeking a deeper explanation of the DC dynamics from the chemical-mechanical coupling, but they have incorporated much phenomenology and left several fundamental questions unanswered. What is the biochemical mechanism for the spontaneous oscillation in the early phase? What mechanism organizes the ratcheting behavior in the early phase? What triggers the transition from one phase to the next?

These questions have motivated the current model, which integrates earlier vertex models (1, 17) with the biochemical insights of recent experiments (4, 12) to provide a comprehensive picture of dorsal closure. In particular, we will demonstrate how the PAR proteins interact with a regulator of the actomyosin network to produce not only the oscillation in the early phase, but also the ratcheting in the slow phase and the persistent contraction in the fast phase. Essentially all features of the DC process, except for apoptosis of AS cells and zippering that affect the final closure, can be explained by a cell-mechanical framework supplied with a few experimental observations regarding the movement and biochemistry of the PAR proteins.

## MODEL FORMULATION

We represent the amnioserosa tissue by a 2D vertex model of 121 hexagonal cells (Fig. 1a). Since the AS cells are thin and squamous, and the actomyosin and signaling proteins of interest are localized to the apical surface, we adopt a 2D planar representation (17). Each cell has six peripheral nodes connected by passively elastic edges due to the actin cortex. A central node is connected to the circumference by elastic spokes that also have an active force component due to myosin contraction (Fig. 1a inset). The amnioserosa is flanked by an epidermis that resists dorsal closure (16, 31), but can also engage in active elongation (24). As a simplified treatment, we surround the amnioserosa by elastic tension lines (blue lines in Fig. 1a) attached to an outer boundary that is fixed in space. The mechanical model essentially follows the previous work of Wang et al. (17). More details, including the treatment of the supracellular actin cable and the epidermal tension, can be found in the online Supporting Material (SM).

**Figure 1:**
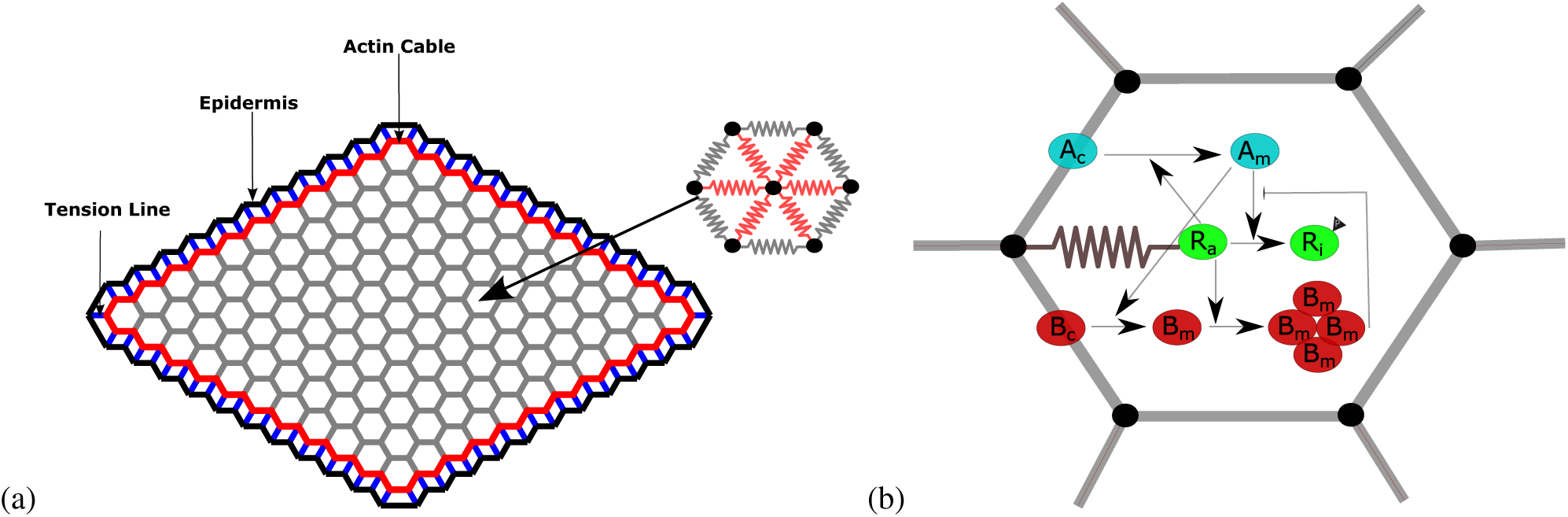
(a) The AS tissue comprises 121 hexagonal cells and the big arrow points to a “representative cell” whose dynamics is reported later in the paper. Each cell has 6 edges (grey) and 6 spokes (red). The edges and spokes are passively elastic represented by springs and the spokes also carry an active myosin force. The AS tissue is linked by blue elastic tension lines to a stationary epidermis (black outline), and the magenta border represents the supracellular actin cable. (b) The translocation and interactions of the PAR proteins inside each cell. Apicomedial actomyosin, represented by its regulator *R_a_*, recruits apico-circumferential aPKC (*A_c_ → A_m_*) medially to form a pool of apicomedial aPKC, which phosphorylates the active regulator *R_a_* into an inactive form *R_i_*. Concomitantly, *A_m_* recruits its own inhibitor Baz medially (*B_c_ → B_m_*), which in a clustered form sequesters *A_m_*. *R_a_* promotes the clustering of *B_m_*.

Our kinetic model is novel in its detailed treatment of the molecular signaling pathways, based on observations of the transport and interaction of the PAR proteins (4, 12, 32), the identity of the target protein for aPKC (33–36), and the clustering dynamics of apicomedial Baz (37–39). Figure 1(b) depicts the protein species tracked in our model as well as their spatial localization and interactions. The PAR proteins, Baz, Par-6 and aPKC, have been known to regulate the apicomedial localization and activation of actomyosin in amnioserosa (4, 12). Of the three, PAR-6 always acts in conjunction with aPKC (4, 40), and we will consider the aPKC-Par-6 complex a single entity in the following, referring to it simply as aPKC. Although aPKC is long known to inhibit actomyosin, its target, which is presumably a regulator or effector of RhoA/Rho1, has not been unequivocally identified. In mammalian cells, Smurf1 can be phosphorylated by aPKC to cause RhoA degradation (33), and aPKC may also act through p190RhoGAP to inhibit RhoA (34). More recent evidence points to Rho kinase (Rok) as a likely target, as phosphorylation by aPKC causes Rok to dissociate from the cell cortex (35). In *Drosophila* placode border, aPKC phosphorylates Rok to deactivate its role in recruiting myosin (36). Based on these, Rok is a strong candidate as the myosin regulator during DC. To be prudent, however, we will use a generic name Reg in the model.

Another important feature of Fig. 1(b) is the localization and translocation of aPKC and Baz. At the onset of DC, the kinase-active store of aPKC is mostly bound to the plasma membrane to either of the two membrane proteins Cdc42 or Crumbs (41). As aPKC is prevented from accumulating on the basolateral plasma membranes by a molecular exclusion mechanism (41), it has been observed mostly on the apical surface, and concentrated along the circumference (4, 12). Meanwhile, Baz is also concentrated in the adherens junctions in the apico-circumference region (41). During DC, the circumferential aPKC and Baz are progressively recruited to the apicomedial domain (12). More specifically, aPKC is recruited by the actomyosin network, and it in turn recruits Baz. Both proteins form puncta in the apicomedial region of the cell. Schematically, Fig. 1(b) uses two arrows to represent these two successive recruitments:

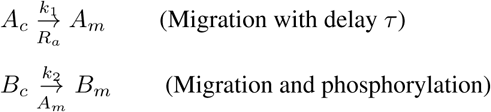

where we indicate both the rate constant and the recruiting agent for each migration. The exact mechanisms of the circumference-to-apicomedial migration are unknown at present, although several possibilities, e.g. diffusion through the cytosol, direct translocation of large protein complexes along the membrane, and actomyosin advection, have been suggested based on *C. elegans* data (37–39). To account for the multiple steps involved in this process that are not explicitly modeled, we introduce a time delay τ in converting the circumferential aPKC *A_c_* to the medial aPKC *A_m_*. For the transport of Baz, on the other hand, we do not implement an explicit delay for several reasons. The recruitment of *B_c_* by *A_m_* represents an interaction that is quickly reversed by aPKC phosphorylation of Baz (12). Besides, as explained below, *B_m_* undergoes an oligomerization into a clustered form before its interaction with *A_m_*. This two-step process will be modeled explicitly, with kinetic rates that amount to an effective delay that allows *A_m_* the time to inactivate *R_a_*. Finally, numerical experimentation shows that an explicit delay in Baz transport is non-essential and does not affect the qualitative outcome of the model.

Once medial, the PAR-proteins undergo a series of dynamic interactions that eventually leads to the cyclic assembly and disassembly of the medial actomyosin network in each AS cell. Medial aPKC inhibits contraction by binding with an active form of Reg (*R_a_*) followed by a phosphorylation event and a quick unbinding, yielding an inactive form of Reg (*R_i_*) and the original aPKC molecule (35, 36):

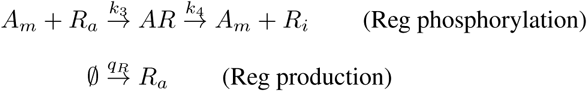

We neglect specific reactivation of the inactive form of Reg (*R_i_*) into the active form *R_a_,* and instead account for such reactivation through a Reg source *q_R_*. This is motivated on the one hand by observations that an expression of upstream actomyosin regulators, e.g. DRhoGEF2 acting on Rho1 for downstream activation of Rok or Diaphonous (13), is necessary to maintain wild-type AS cell behavior. On the other hand, *q_R_* also enables the model to predict a rising level of apicomedial myosin in later stages of DC, which is a prominent feature *in vivo,* both at the protein level (4, 12) and at the level of induced gene expression in the tissue (42).

The apicomedial Baz molecules (*B_m_*) become clustered 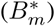 under the influence of *R_a_*. Imaging evidence suggests that this occurs through the mechanical tension produced by the networks rather than chemical binding to the actomyosin; the tension may enhance Baz oligomerization through structural reorganization of the multi-domain protein (37, 43). The Baz clusters 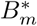 then sequester *A_m_* and antagonize its effect on myosin regulation (12, 37–39):

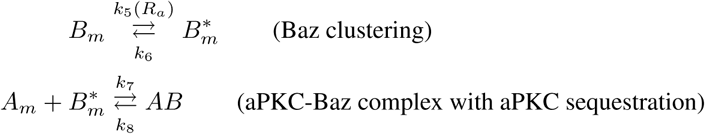

These reactions are indicated respectively by pointed and blunted arrows in Fig. 1(b). The sequestration of aPKC by Baz clusters has an uncertain stoichiometry. It is known that each Baz protein has multiple binding sites for the Par-6-aPKC complex (12, 44). Depending on the number of Baz in the cluster, each cluster can potentially sequester a large number of aPKC molecules before it is saturated. For lack of quantitative knowledge of the stoichiometry, we have made the simplifying assumption that a Baz cluster, once formed, can continue to sequester aPKC without being saturated. As will be seen later, this affects the model only in one minor aspect: the consumption of Baz in the medial domain.

The interactions described above establish a competition for *A_m_* by *R_a_* and 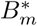, and a potential scheme for generating a delayed negative feedback loop as hypothesized by David et al. (12). The arrival of *A_m_* at the apicomedial domain antagonizes *R_a_* and causes the actomyosin network to disassemble. But *A_m_* also recruits *B_m_*, whose arrival and clustering yield an agent that inhibits *A_m_*, giving *R_a_* the time to accumulate and reassemble the actomyosin network. The inhibition of *A_m_* by 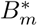 is helped by the fact that the dissociation rates *k*_6_ for 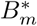 and *k*_8_ for *AB* are both slow, making 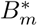 a potent aPKC sequestration mechanism (37–39). The transport and kinetic processes can be described by the following coupled differential equations, all protein species being expressed in terms of concentration:

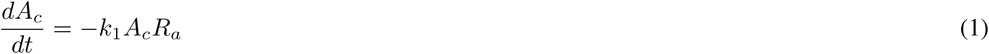

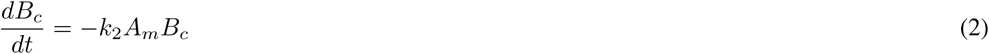

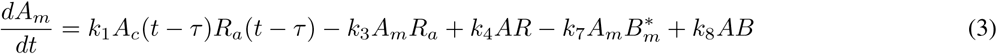

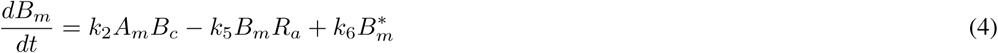

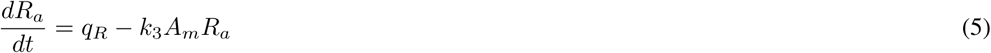

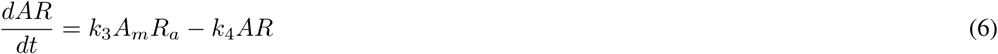

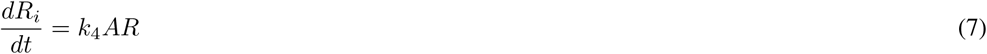

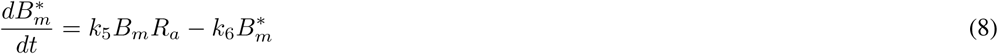

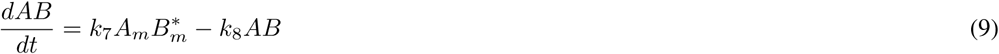

The regulator *R_a_* serves as the link between the kinetics of the biochemistry and the mechanics of actomyosin contraction. The scheme of distributing *R_a_* in each cell among the six spokes and the kinetic equation for the attachment and detachment of myosin motors are the same as in Wang et al. (17), and details are given in the SM. The model has a number of parameters, which are evaluated from experimental data where possible. To address uncertainties in the parameter values, we have performed a sensitivity analysis and found the kinetic model to be robust around the chosen values. Details of this analysis, along with a table of the parameter values, can be found in the SM.

We start the simulation with an initial circumferential aPKC store of *A_c_* and Baz store of *B_c_*, and no Reg or myosin in the medial region. In the real amnioserosa, the cells exhibit phase lags among the neighbors and they oscillate preferentially in antiphase (1). To introduce the phase lag, we stagger the start of the kinetic model among the cells by a time randomly chosen between *t* = 0 and *t* = *T*, where *T* ~ 260 s is the period of oscillations observed in experiments (1) and also captured by our simulation.

## RESULTS

Our model is able to predict the three distinct phases of dorsal closure. Figure 2 depicts the temporal evolution of the area of a representative cell in panel (a) and that of the whole tissue in panel (b). The early phase features an oscillating tissue area without a sustained decrease over the cycles. This gives way to the slow phase that starts with the onset of net tissue area loss and ends with the cessation of oscillation on the cell level. The total loss in tissue area amounts to about 18% during the slow phase. Finally, the fast phase is distinguished by mostly monotonic shrinking of cell area and a rapid contraction of the tissue area toward closure. The dynamics of the entire process can be viewed in Movie S1 of the online SM. In the following we explore the molecular mechanisms that underlie each phase and trigger the transition from one to the next.

**Figure 2:**
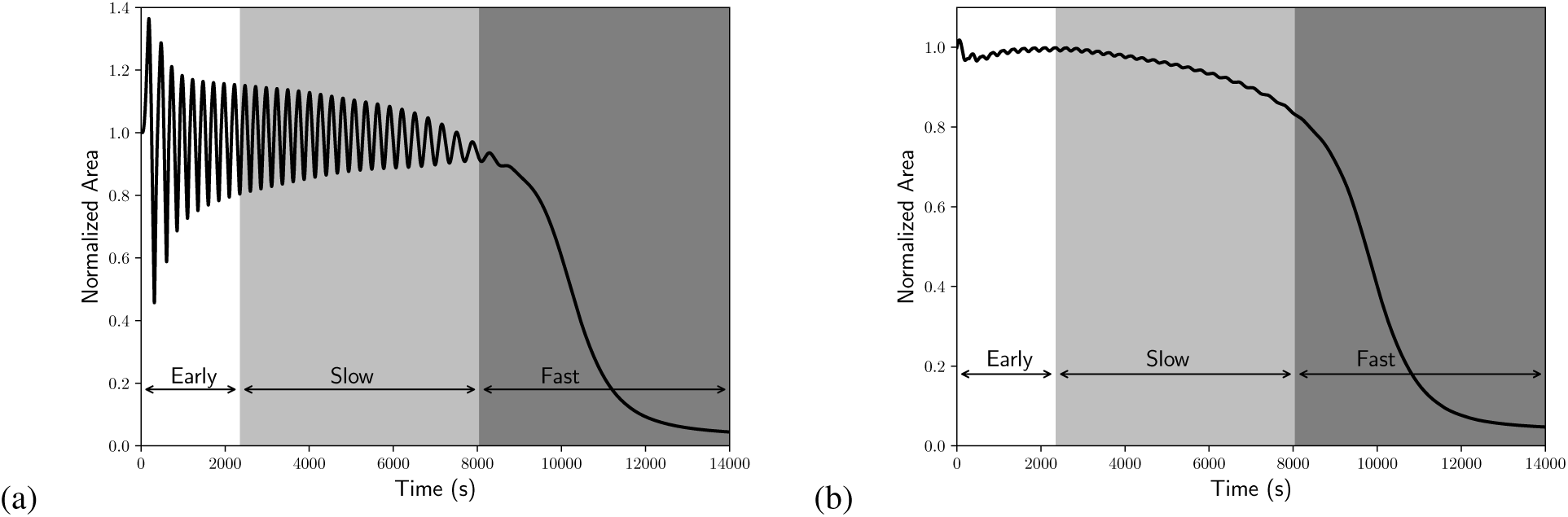
The temporal evolution of (a) the area of the representative cell of Fig. 1 and (b) the entire tissue area. Both areas are normalized by their initial values. The early phase shows oscillatory behavior, the slow phase shows dampening oscillations and gradual area loss, and the fast phase shows persistent loss of area.

### Early Phase: Oscillations

The purpose of this subsection is to investigate the origin of the oscillatory behavior of the AS cells within the context of the chemical signaling pathways detailed in the kinetic model. Figure 3 shows the early-phase dynamics of the myosin motors and cell area of the representative cell highlighted in Fig. 1(a). For this cell, the PAR-protein network begins at *t* = 179 s, before which the cell is only subject to mechanical forces from its neighbors. Once the biochemical network is activated, cell behavior becomes dominated by the internal myosin dynamics. Persistent oscillations of the cell area and the myosin level develop in time. The mean period of oscillations *T* ≈ 260 s matches experimental observations (1, 3). Parametric studies show that *T* is most sensitive to the delay parameter *τ* (see SM), and *T* increases with *τ*. This suggests that the period of oscillations is an outcome of the transport and activation mechanisms of apicomedial aPKC. Moreover, the cell area oscillates roughly in anti-phase with the myosin, with an amplitude of about 20% among AS cells (Fig. 3). The maximum of myosin precedes the minimum of the cell area slightly, in agreement with experimental observations (3, 4, 17). We have also calculated the cross correlation of the area change between neighboring cells. Although the cells have been given a random initial phase lag, once they settle into a regular oscillation, the mechanical coupling among neighbors produce on average a negative correlation (Fig. S6 in SM), again in agreement with experimental observations (1, 3).

**Figure 3:**
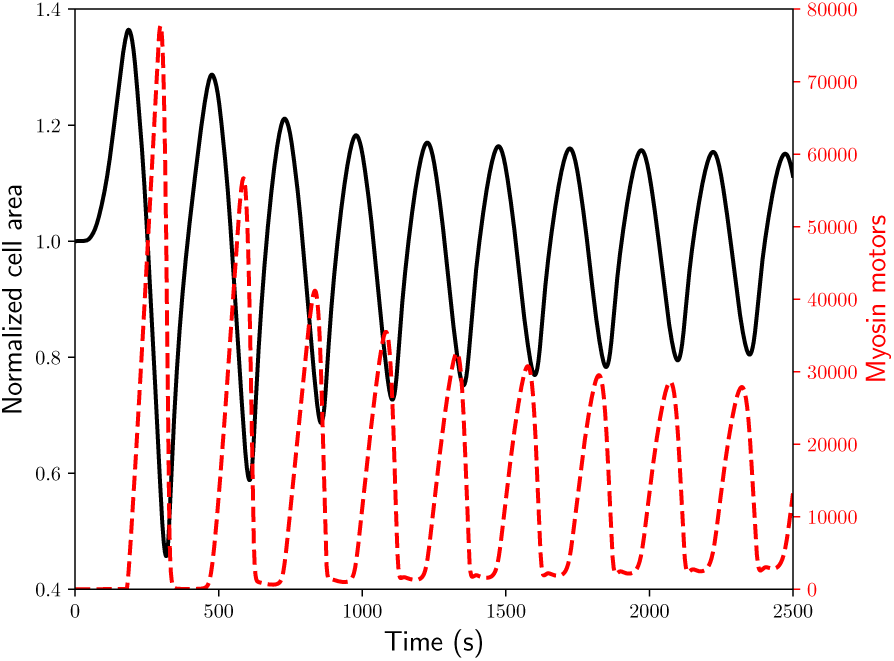
Early phase dynamics of the normalized area of the representative cell (black solid line) and the total myosin inside the cell (red dashed line).

The area of the entire tissue also exhibits temporal variations (Fig. 2b, Movie S1), but the overall change is less than 6% during the early phase. As the actomyosin network is activated throughout the AS cells, the tissue area first declines by about 5% and then recovers to roughly its initial area. The decline is owing to the fact that the first few cycles of myosin oscillation are particularly strong, with large amplitudes (cf. Fig. 4a below). In time, it settles into a milder oscillatory pattern, producing the recovery in tissue area. This process manifests itself in the evolution of individual cell areas by a decrease in the amplitude of the area (Fig. 2a). One can also discern small-amplitude oscillations in the whole tissue area, a collective effect of the oscillation of all the AS cells. In addition, we see no difference in behavior between the leading edge cells and those in the interior.

**Figure 4:**
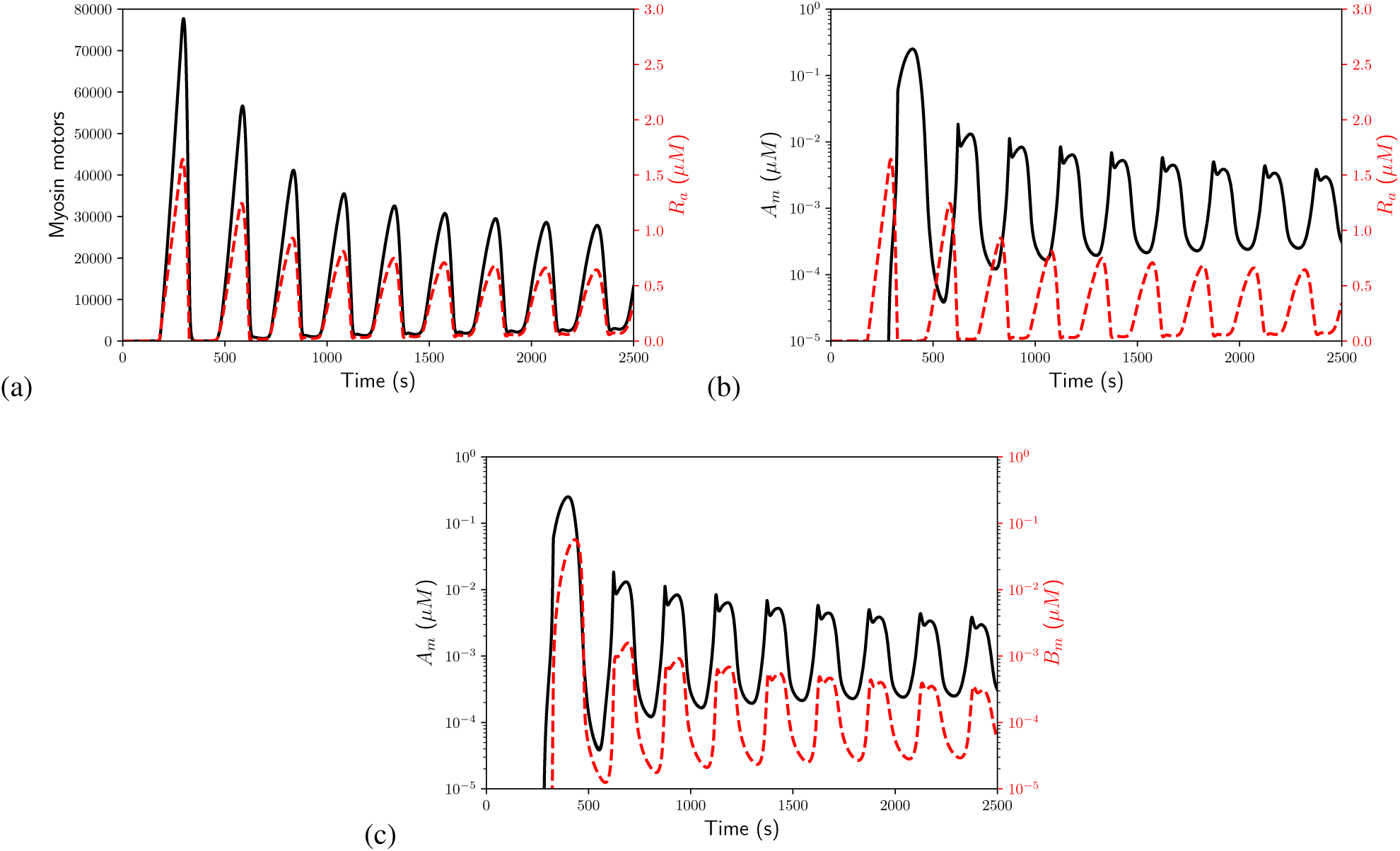
Evolution of various medial proteins in the representative cell during the early phase. (a) Number of myosin motors (black) and concentration of the actomyosin regulator *R_a_* (red dashed). (b) *R_a_* (red dashed) and its antagonizer aPKC *A_m_* (black). (c) *A_m_* (black) and Baz *B_m_* (red dashed).

The origin of the oscillation can be sought from temporal and spatial relationship among the PAR proteins and the regulator. The direct cause for the myosin pulse is a pulse in *R_a_* (Fig. 4a). *R_a_* grows because of the steady production *q_R_* (Eq. 5), and its growth recruits *A_m_* with a delay *τ* (Eq. 3). As *A_m_* accumulates in the apicomedial domain, it continues to phosphorylate and deactivate *R_a_*, causing the latter to peak and then sharply decline (Fig. 4b). Thus, the medial myosin network disassembles as a result of the medial aPKC (4). Meanwhile, *A_m_* recruits Baz (Eq. 4) so *B_m_* rises in tandem with *A_m_* (Fig. 4c). As *B_m_* oligomerizes into 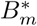 that in turn sequesters *A_m_, A_m_* declines. With the now depressed level of *A_m_*, *R_a_* is able to recover and re-assemble the actomyosin network, which sets up the next cycle of oscillation. The duration of the actomyosin contraction depends on how long *A_m_* persists with Baz. Higher Baz prolongs the contraction phase as observed by David et al. (4).

It is interesting to note that the amplitude of the oscillation gradually declines in time for all proteins. The first few cycles have particularly large amplitudes. This is related to our model assumptions that initially all aPKC and Baz are stored in the circumference of the cell’s apical surface, and that the rate of recruitment to the medial domain is proportional to the entire stores of *A_c_* and *B_c_* (Eqs. 1, 2). Thus, the first few large waves are incidental to the setup of the simulation and practically inconsequential. However, a more consequential feature is that the circumferential stores of aPKC and Baz are finite and decline in time as more of them translocate to the apicomedial domain to mediate the actomyosin dynamics. We will return to the temporal decline of the amplitudes when discussing the transition to the slow and fast phases.

Thus, the model explains the oscillatory behavior of the AS cells from the *cell-autonomous* kinetics of signaling proteins involved in DC. The question of whether cell oscillation arises from intracellular or intercellular mechanisms has been debated in the literature. Some mathematical models, including the present model and that of Machado et al. (8), can produce oscillation from cell-autonomous mechanisms, while others rely on mechanical coupling between neighbors (1, 17). There is little doubt that the contractile phase of the oscillation is due to myosin motors inside each cell. It is the expansion phase that is in question. Experiments using holographic laser microsurgery (27) suggest that cell expansion is driven by intracellular forces, and is thus cell-autonomous. More recent data (9) paint a subtler picture in which a combination of autonomous and non-autonomous mechanisms drive cell expansion during AS oscillation. In our model, expansion occurs via passive elasticity of the cell, whose structural basis must be the actomyosin cortex. In this sense, our model is consistent with the cell-autonomous expansion noted by Hara et al. (9). But it does not discount an alternative pathway to oscillation through cell-cell coupling, which may coexist and cooperate with the cell-autonomous pathway in the real amnioserosa.

To sum up this subsection, our model is closely based on the observed intracellular kinetics of PAR proteins during DC (4, 12) and their effects on actomyosin observed in *Drosophila* and other organisms (35, 36). The possibility of aPKC phosphorylating Reg and complexing Baz results in a competition between the latter two, and sets up negative feedback from aPKC onto actomyosin and from Baz onto aPKC. Hence the oscillation arises. By encoding the biological insights into a quantitative form, the model demonstrates a cell-autonomous mechanism for the oscillation.

### Slow Phase: Ratcheting

The beginning of the slow phase is defined as the onset of net contraction of the AS tissue (2, 3). In Fig. 2(b), we identify this as the instant *t* = 2352 s when the tissue area reaches a maximum before starting to decline. The end of the slow phase is marked by the cessation of cell area oscillation and a dramatic acceleration of AS contraction rate (3). Using the data of Blanchard et al. (3) as a reference and considering the variation among our model cells, we have determined the end of the slow phase at *t* = 8046 s, giving the slow phase a duration of 5694 s or roughly 95 min. Overall the slow phase is characterized by a gradual intensification of net area contraction. Measured by its maximum area in each cycle, our representative cell of Fig. 2(a) loses about 0.5% of area per cycle at the start, up to 3% per cycle toward the end, for a cumulative reduction of 20% during the slow phase. There is also a dampening of the amplitude of areal oscillation; it decreases from roughly 20% of the mean area to about 2% for the representative cell.

Our model captures the net decrease of cell area through an increasing average myosin level. This is evident from the temporal evolutions of both in the slow phase (Fig. 5a). With each cell cycle, the minimum myosin level increases (red arrow); it stabilizes the cell cortex and prevents the cell from relaxing to its original area (black arrow). Tracing the biochemistry upstream, the increasing myosin level is the result of a gradually rising *R_a_*, which in turn is due to its source *q_R_* and a decline in the level of apicomedial aPKC *A_m_* (Fig. 5b). As we have started with a finite reservoir of circumferential aPKC, *A_c_*, the total amount of *A_m_* is also finite. As more of it is sequestered by 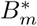, there is progressively less available *A_m_*. As a result, *R_a_* rises over each cycle.

**Figure 5:**
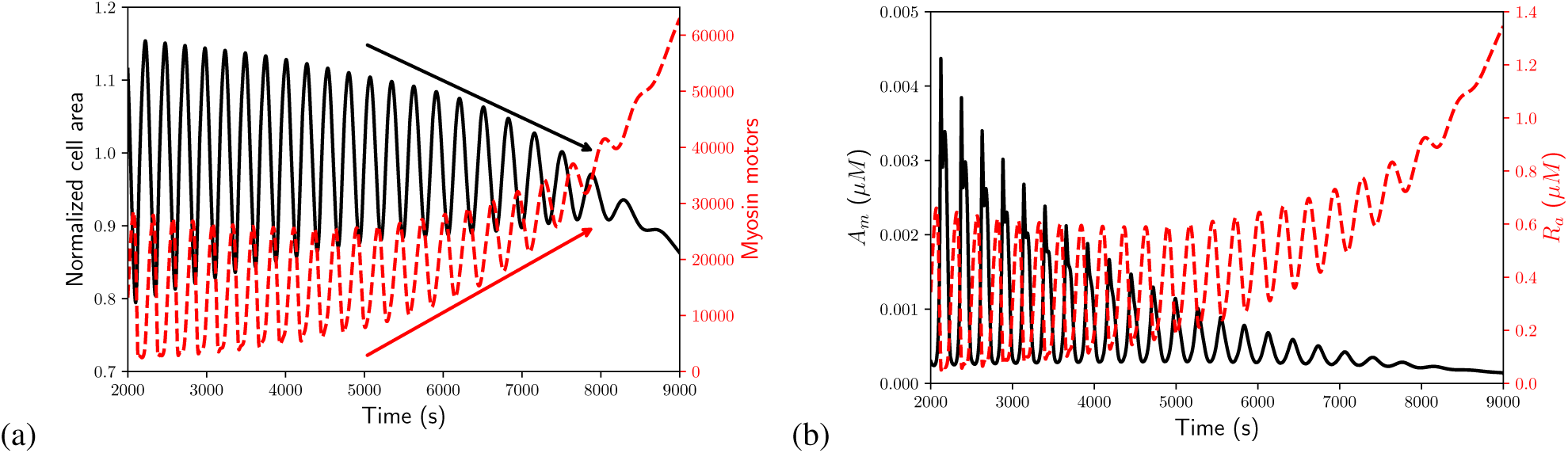
Evolution of various proteins in the representative cell during the slow phase. (a) Over each cycle of oscillation, an increase in myosin (red dashed) leads to constriction of the cell area (black), resulting in a ratchet-like effect. (b) The active Reg *R_a_* (red dashed) rises over each cycle as the medial aPKC *A_m_* (black) declines.

A central concept in the current view of cell constriction is the internal ratchet (7). However, there is no convincing model or explanation of its underlying mechanism. The current hypothesis is that the cell cortex and internal actomyosin network stabilize themselves at the conclusion of every myosin pulse, thus preventing the cell area from returning to the previous value before the contraction (1, 7, 11, 15). Then what stabilizes the actomyosin after each contractile pulse? Why is this behavior unique to the slow phase of DC, but not present in the oscillatory early phase? In their DC model, Wang et al. (17) circumvented these questions by prescribing an *ad hoc* “mechanical ratchet”, reducing the resting length of each cell edge and spoke by a prescribed amount during the slow and late phases. Our model answers these questions on the basis of the biochemistry of the PAR proteins. During the early phase, there is still sufficient aPKC to antagonize the actomyosin regulator and allow the actomyosin network to disassemble completely. Consequently, the cell relaxes to its original area. The early phase comes to an end with the gradual exhaustion of apicomedial aPKC. In the slow phase, the declining level of aPKC is such that it is no longer able to completely suppress net myosin production in each cycle, resulting in a cumulative increase in the number of myosin motors with each successive cycle. This, our model suggests, is the molecular basis for the function of the actomyosin ratcheting.

The area of the entire AS tissue also shrinks gradually (Fig. 2b), the contraction amounting to roughly 18% by the end of the slow phase. Experimentally, the beginning of the slow phase has been observed to coincide with the formation of the supracellular actin cable surrounding the AS tissue (2, 3), suggestion a major role for the actin cable in AS contraction (1–3). Follow-up studies concluded, however, that the actin cable is not the main cause of closure (6, 18, 19), and contributes only a small fraction of the total area loss (17). Considering its relatively minor role, we have chosen to model the actin cable *ad hoc* at a coarser level than the intracellular dynamics (details in SM). Our model predictions confirm the insignificance of the actin cable to dorsal closure. Turning on the actin cable at a prescribed time causes a slight decrease in the tissue area, relative to the case without actin cable (Fig. S7a), but the effect is minimal (up to 2%). Visually, the actin cable tends to align the outermost cell edges so as to give the tissue a smoother leading edge. This is similar to the prediction of the earlier model of Wang et al. (17). In addition, our results confirm that the actin cable is not the trigger for the start of the slow phase. When we activate the actin cable at different times, an early onset does produce a smaller tissue area throughout, but the difference is minuscule (Fig. S7b). In particular, the instant at which the tissue area peaks before declining, which is taken to be the start of the slow phase, is not affected by the actin cable. Movies S2 depicts the entire closure process with the actin cable deactivated.

We close this subsection by noting two experimental features that the model fails to reproduce. First, Solon et al. (1) reported that the cessation of cell oscillation occurs first at the boundary with the epidermis, and propagates inward toward the midline of the AS. In our model, all cells display roughly the same amplitudes of oscillation as the PAR-protein dynamics are largely cell-autonomous. Thus, the cessation of oscillation occurs randomly throughout the tissue, depending on the initial phase lag among the cells and their mechanical coupling. Second, *in vivo* the amplitude of the oscillation dampens and its period shortens as the slow phase progresses (3). Our model captures the dampening of the oscillation due to the gradual buildup of apicomedial myosin, but not the reduction in the period *T*. Our period actually lengthens, from about 260 s at the start to 350 s toward the end (Fig. 2a). Evidently, our mathematical description of the PAR-proteins misses some subtle mechanism that shortens the oscillation period *in vivo.* In the model of Machado et al. (8), AS cell oscillation arises from the cyclic actin turnover, and these authors were able to produce a shortening period by prescribing an actin turnover rate that increases in time. But at present it is unclear what upstream mechanism could accelerate the actin turnover rate.

### Fast Phase: Closure

The fast phase, as observed *in vivo,* introduces two new factors into the closure of the AS tissue. The first is zippering (20–22), which occurs when filopodia are extended from one epidermis to the other and exert a pulling force to move the two leading edges toward each other. Filopodia are approximately 10 μm in length, and function only after the internal actomyosin contraction has brought the leading edges to within one or two cell widths. We have chosen to exclude this factor from the model because it is non-essential for dorsal closure (6) and is largely independent of the AS contraction. The second is apoptosis and delamination of AS cells (2, 5, 23), which mainly occurs at the canthi during the onset of the zippering process and contributes to the speed of dorsal closure. We model this process by eliminating a cell when its area falls below a threshold area (1 μm^2^) and all its edges are shorter than 0.5 μm. To maintain the integrity of the model tissue, we then collapse the outer nodes of the “apoptotic” cell onto the central node, a treatment motivated by *in vivo* experiments of Toyama et al. (23) and their Figure 2(a,b). If eliminating a cell brings the opposing epidermis together, we consider this the fusion of the epidermis and subsequently fix the location of the central node. This mimics the restructuring process that occurs at the canthi *in vivo* (6).

Upon the onset of the fast phase, the AS cells stop oscillating and undergo persistent constriction while the tissue area drops precipitously (Fig. 2b). The rapid loss of AS area lasts about 4000 s, as the rate of contraction slows down. It takes roughly another 2000 s of slow contraction to reduce the AS areas to below 5% of its initial value. This is considered “full closure” in our model, considering the lack of zippering. This duration of the fast phase is consistent with experimental observations (5, 24). Movies S1 in the online SM gives an overall view of the whole process.

The evolving geometry of the amnioserosa is depicted in Fig. 6. At the start of the fast phase (*t* = 8046 s), the geometry of the AS tissue resembles that of a rounded rhombus (Fig. 6a). As contraction becomes more intense, the outermost cells on the anterior-posterior axis are the first to become eligible for apoptosis (Fig. 6b). Their elimination forms two canthi around *t* = 10100 s. Once the canthi fix the length of the AS tissue, it narrows rapidly into a spindle shape (Fig. 6c) that closely resembles the *in vivo* observation (6). In time, our AS tissue is able to shrink its area to below 5% of the initial value, producing a thin slit between the two epidermal tissues (Fig. 6d). This is the final “fully closed” geometry. Closure is a robust prediction of our model that is attained with and without the actin cable (Fig. S7), and over wide ranges of the kinetic parameters (see parametric study in SM).

**Figure 6:**
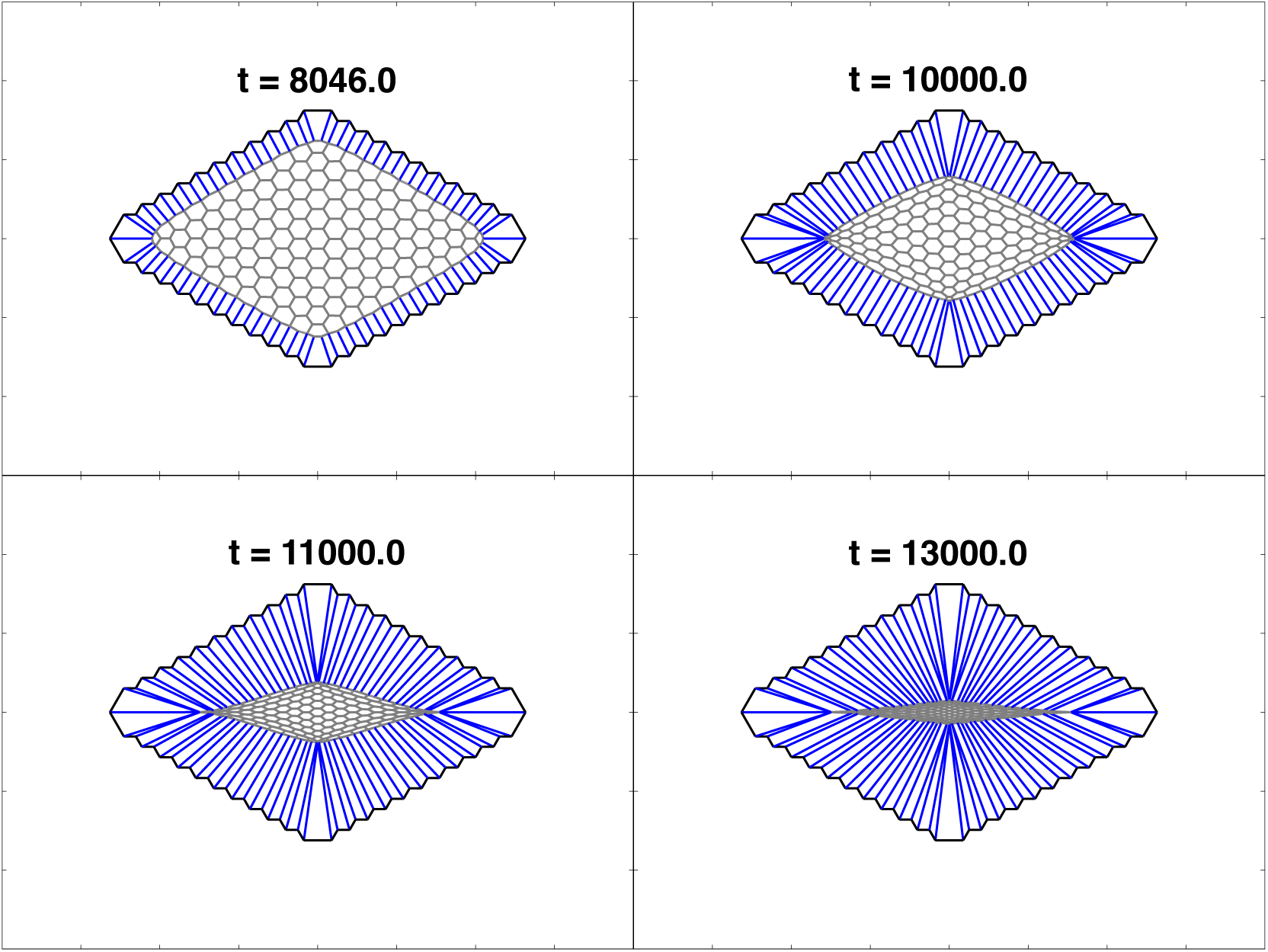
Contraction of the amnioserosa in the fast phase. (a) At the start of the fast phase *t* = 8046 s, the tissue boundary has been smoothed by the actin cable as compared to the initial configuration, (b) Shortly before the canthi form around *t* = 10100 s. (c) After canthi formation, the AS narrows rapidly in the medial-lateral direction, (d) At *t =* 13000 s, the tissue area has fallen below 5% of its initial value, and this is considered full closure in the context of our modeling.

What triggers the transition from the slow to the fast phase is the exhaustion of apicomedial aPKC. Through the cycles of actomyosin assembly and disassembly of the early and slow phases, most of the circumferential aPKC (*A_c_*) has migrated to the medial domain, and the medial aPKC (*A_m_*) is gradually sequestered by 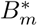. Figure 7(a) shows the declining amplitude and mean of *A_m_* in the early and slow phases, as each cycle receives less influx of aPKC from the circumferential *A_c_*. Since *B_m_* is recruited by *A_m_* to the apicomedial domain, it follows a similar trend of decline. By the end of the slow phase, *A_m_* and *B_m_* have more or less approached a steady state, as do the products *AB* and *AR* (Fig. 7b). Henceforth, *A_m_* and *B_m_* have essentially dropped out of the negative feedback that antagonizes *R_a_* and suppresses the assembly of myosin. This triggers the transition into the fast phase as the myosin pulsation gives way to a monotonic rise (Fig. 7c), and the cell-area oscillation to persistent constriction. Note that the steady-state *B_m_* concentration is considerably lower than that of *A_m_*. This observation is consistent with experimental observations of less Baz than Par-6 in the apicomedial complexes (12).

**Figure 7:**
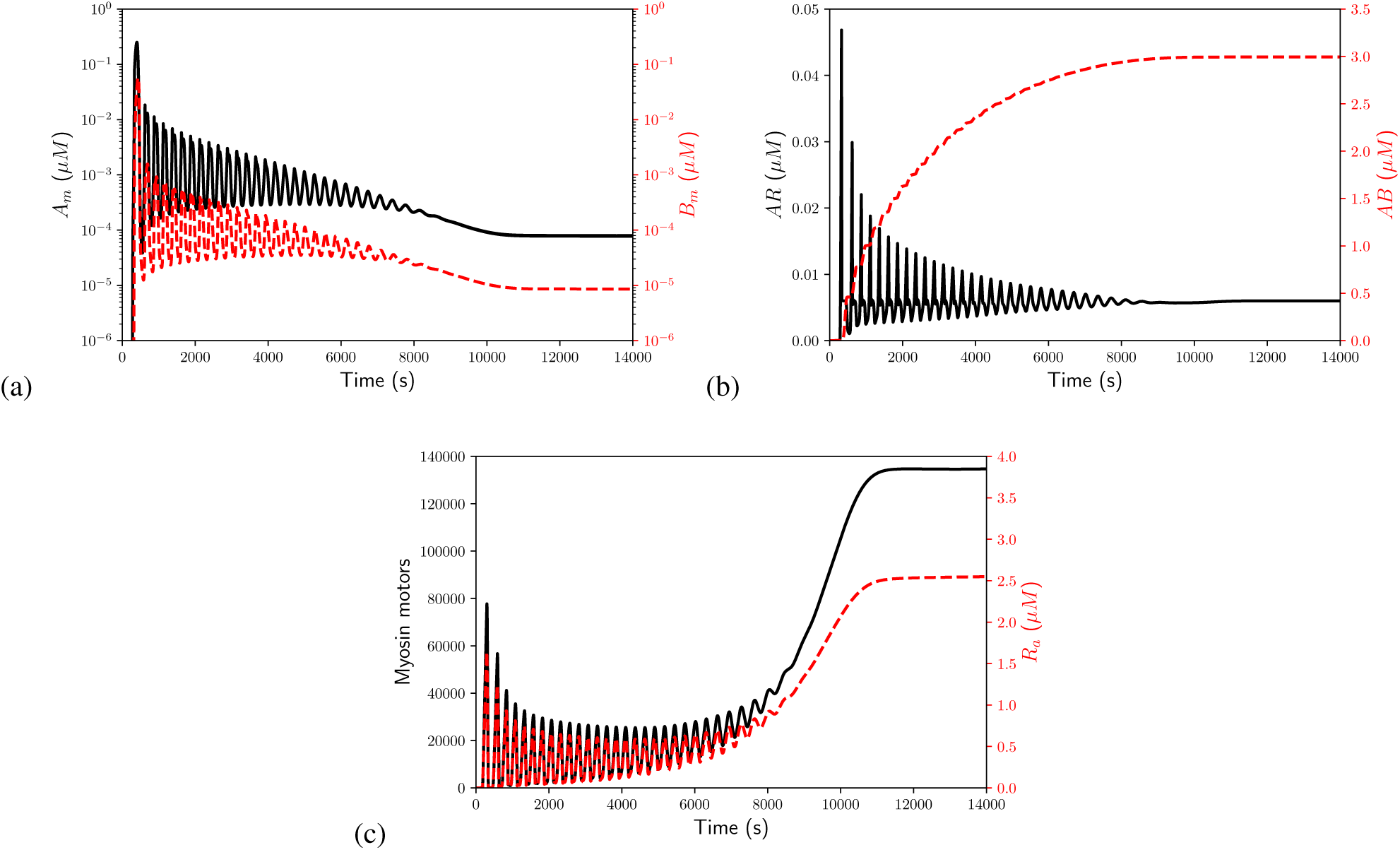
Dynamics of the key proteins in the representative cell explain the transition into the fast phase. (a) *A_m_* (black) and *B_m_* (red dashed) decline, as medial aPKC is consumed by phosphorylation of Reg and Baz. (b) The products *AR* (black) and *AB* (red dashed) attain a steady state in time. (c) Oscillations in myosin (black) and *R_a_* (red dashed) in the slow phase give way to monotonic growth in the fast phase.

Interestingly, we find that the circumferential Baz store *B_c_* has little effect on the onset of the fast phase. While the entire circumferential store *A_c_* is eventually recruited by Reg to the apicomedial region, the same is not true for *B_c_* (Fig. S8). As *A_m_* declines in the fast phase (Fig. 7a), it can no longer recruit *B_c_* effectively. Thus, the transport of *B_c_* to *B_m_* dies out, with only about 4% of *B_c_* having migrated to the apicomedial domain. Such a low level of apicomedial Baz is sufficient to carry out its task of sequestering aPKC, partly because of our assumption of 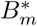 not being saturated in its capability of sequestering aPKC (Eq. 8). Biologically, the rationale is that the Baz cluster 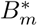 has multiple binding sites for aPKC, and thus is not rendered ineffectual after forming the first complexes with aPKC (12, 32, 44, 45). Nevertheless, the large initial store of *B_c_* is necessary for the early and slow phase. Our kinetics is such that the rate of Baz migration is proportional to *B_c_*. Thus, the large *B_c_* maintains a sufficient rate of *B_m_* that is needed for suppressing aPKC and allowing the actomyosin to reassemble.

Experimental consensus is that in the fast phase the PAR-proteins progressively accumulate apicomedially and the acto-myosin network becomes more persistent and pervasive throughout the AS tissue (12, 21). These observations guided the construction of the mathematical model described in Eqs. (1–9), and are recapitulated by our simulation results. The strengthened actomyosin network dampens the AS cell oscillations and effects persistent constriction (1, 3). Our model has predicted a natural transition from the slow to the fast phase, and also captured its key features noted above. In this regard, it marks an advance beyond previous modeling, which has rarely dealt with the fast phase (1, 8, 27, 28). The model of Wang et al. (17), for example, was unable to predict the upsurge of apicomedial myosin in the fast phase. There was no chemical or mechanical cue to trigger the transition from oscillatory to non-oscillatory dynamics in the apicomedial myosin dynamics. In its stead, a mechanical ratchet was employed, by reducing the rest length of the edges and spokes in time, to produce rapid closure of the tissue.

## DISCUSSION

The objective of this work is to integrate experimental findings of the molecular signaling pathways in *Drosophila* dorsal closure into a quantitative and predictive model. The model has two key components. One is a PAR-protein regulatory circuit that has been suggested by recent observations (12, 32, 37–39). The other is a vertex-based cell mechanics model that is driven by myosin contraction. The myosin activation by a regulator downstream of the PAR-kinetics provides the linkage between the two. The model parameters have been evaluated based on available data, and the robustness of the results has been tested by varying these parameters. The main results of the paper consist of the following:

- The model predicts the oscillations in the apical area of amnioserosa cells in the early phase, and provides an explanation for it from the cell-autonomous biochemistry of the PAR proteins.
- The model predicts the internal ratchet that causes gradual loss of cell area in the slow phase of dorsal closure, and explains its molecular mechanism based on the same biochemistry.
- The model predicts the cessation of cellular oscillation in the fast and final phase of dorsal closure, as well as the rapid and persistent contraction of cell and tissue area, again from the biochemistry.

The model predictions are in reasonable agreement with experiments. For example, it reproduces the various time scales of the process, including the period of oscillation and the duration of the three phases, confirms the anti-phase correlation among neighbors and the minor role of the supracellular actin cable, and recapitulates the temporal dynamics of the key proteins, especially the intensification of apicomedial myosin in the later stage of dorsal closure. Therefore, from a minimal regulatory circuit and cell mechanics, the model is able to capture most key features of the experimental observations in the early, slow and fast phases, as well as the transition among them. It provides a coherent framework that connects the tissue- and cell-level events to the intracellular transport and interaction of the PAR proteins. These are the main contributions of the current work.

Of the factors contributing to dorsal closure, we have accounted for the intracellular contraction of actomyosin, the supracellular actin cable, the interaction with the surrounding epidermis, and the delamination of apoptotic cells. Zippering, an important mechanism during the final stage of closure, has been neglected completely. Of the factors included, the actomyosin contraction is modeled with the greatest care; it is the centerpiece of the model. The three other factors are modeled more *ad hoc,* using convenient mechanical and mathematical constructs. This is partly for lack of an understanding of the underlying mechanisms, and partly for their secondary roles in the dorsal closure process. As more is learned about them, the new knowledge can be used to refine the modeling. The model predicts the oscillatory dynamics of the early phase from a cell-autonomous mechanism. This leaves aside, but by no means discounts or disproves, potential roles of cell-cell coupling. Mechanical coupling occurs through forces and deformation, as accounted for by earlier models (1, 17). There may also be biochemical coupling, as evidenced by the intriguing phenomenon of an actomyosin pulse propagating between neighbors (4). This suggests potential pathways through cadherin-based mechaniobiology at cell junctions (46, 47), or through mechanically gated ion channels (48).

Of necessity, the model makes simplifying assumptions about a complex biological process, and omits some of its less important elements. A key assumption is a steady source for the actomyosin regulator *R_a_*, as this lies at the root of the ratcheting and rapid closure in the last two phases. The identity of this regulator has been discussed by several groups (12, 33–36), and Rho kinase appears as a highly plausible candidate among several hypothesized. An obvious task for future experiments is to identify the regulator unambiguously. The assumption of its source and continual supply is also testable in future experiments, by examining its gene expression and medial protein levels in relation to upstream regulators and downstream contractile myosin networks. Besides, for lack of precise knowledge of the stoichiometry of aPKC sequestration by Bazooka clusters, we have assigned unsaturated binding capability to these clusters. An unrealistic consequence is the minimal relocalization of Baz from the circumference to the medial domain, in comparison to *in vivo* observations (e.g. Fig. 2 of David et al. (12)). Future experiments should investigate the complex formation between aPKC and Bazooka and gather quantitative data on the stoichiometry.

The most glaring failure of the model is that it predicts a lengthening period for the AS cell oscillation during the slow phase with internal ratcheting, whereas in reality the damping of the oscillation is accompanied by a shortening of the period (3). We can think of two potential causes for the discrepancy, one geometric and the other mechanical. As DC progresses, the shrinking cellular area should entail shorter migration times for the PAR proteins and potentially accelerate the downstream kinetics. This is unaccounted for in the model. Furthermore, experiments show that the tissue is under increasing tension during DC, and its mechanics changes from fluid-like to solid-like (12, 21, 28). The strengthening of the actomyosin and the rising tension have been captured by the model. However, this gradual rigidification does not translate to an increase in oscillation frequency, as one may expect based on mechanical intuition. The model determines the frequency of AS oscillation mostly from the kinetics of the PAR proteins, and does not take adequate account of the rheological properties of the cells and tissue, including viscoelasticity (28). The various shortcomings of the model may motivate remedies and refinements in future work.

## ACKNOWLEGMENT

We acknowledge financial support by NSERC through Discovery Grant No. 05862 (to JJF) and No. 05617 (to TJCH). We thank William Lou and Amir Mafi for their contribution to an earlier version of the model.

## SUPPORTING CITATIONS

Reference (49–55) appear in the Supporting Material.

